# drugMR: A multi-omics pipeline for genetically anchored drug-target discovery

**DOI:** 10.64898/2026.10.01.756031

**Authors:** Guillermo Comesaña Cimadevila, Marie-Joe Dib, Matthew Bracher-Smith, Fiona Ducotterd, Dervis A. Salih, Nicholas J. Bray, Emily Simmonds, Valentina Escott-Price

## Abstract

**Motivation:** Genetically supported drug targets are an estimated 2.6 times more likely to succeed clinically, yet a disease association alone does not show which protein to target, in which direction, or what other effects may follow. This currently requires chaining separate tools.

**Results:** drugMR is a Nextflow pipeline that, from GWAS and pQTL summary statistics, runs *cis*-Mendelian randomisation, colocalisation, SMR-HEIDI with bulk and/or single-cell QTLs, three-trait colocalisation and a phenome-wide screen for adverse effects and repurposing. Datasets are added through a manifest, every step runs in containers on a laptop, cluster or cloud, and results are explored in a dashboard. Applied to schizophrenia with brain pQTLs, drugMR tested 2,019 proteins in 4 h 19 min and prioritised 58 proteogenomic targets, 22 with multi-omic support; 4 had potential adverse effects.

**Availability and implementation:** drugMR: https://github.com/guillermocomesa_nacimadevila/drugMR; documentation: https://guillermocomesanacimadevila.github.io/drugMR/. Demo data: https://doi.org/10.5281/zenodo.22917384.

## 1 Introduction

Approximately 90% of drug candidates fail after entering clinical development (Sun et al., 2022), and bringing a successful drug to patients can take up to 12 years (Berdigaliyev and Aljofan, 2020). Causal inference using human genetic data can inform target selection before clinical trials, and these genetically supported drug programmes are estimated to be 2.6 times more likely to succeed than those without such support (Minikel et al., 2024).

Mendelian randomisation (MR) is a widely used genetic approach for causal inference. It leverages the random assortment of alleles at conception to estimate causal effects, which is analogous to treatment allocation in a randomised controlled trial (Davies et al., 2018). MR commonly uses genome-wide association study (GWAS) summary statistics, with exposure-associated variants serving as instrumental variables. However, it may be affected by horizontal pleiotropy, whereby a variant influences the outcome independently of the exposure (Verbanck et al., 2018), and analyses of complex traits may not identify a therapeutically actionable protein. This is relevant because approximately 75% of FDA-approved drugs act on specific protein targets (Santos et al., 2016). Protein quantitative trait loci (pQTLs) enable MR of genetically predicted protein abundance, while restricting instruments to *cis*-pQTLs near the encoding gene may reduce, but not eliminate, horizontal pleiotropy (Zheng et al., 2020; Schmidt et al., 2020). Other molecular QTL data, including expression, splicing, methylation and chromatin-accessibility QTLs, can provide supporting evidence across regulatory processes, tissues and cell types, helping to assess whether findings for a target are consistent across multiple omics layers.

Existing summary-data MR pipelines cover only parts of this process. MeRP (Yin and Voight, 2015) automates GWAS-based exposure-outcome MR without colocalisation; Causeway (de Amorim et al., 2025) screens molecular QTLs using MR, pleiotropy checks and coloc; and MRDRP (Mundo et al., 2026) automates *cis*-MR without colocalisation. None combine multi-signal colocalisation, bulk-tissue and single-cell molecular QTL follow-up, and phenome-wide screening for potential adverse effects and repurposing opportunities.

We therefore developed drugMR, a dataset-agnostic pipeline for drug-target prioritisation using multiple omics datasets. Starting with pQTL and outcome GWAS summary statistics, it performs proteome-wide *cis*-MR, single- and multi-signal colocalisation, molecular QTL follow-up using bulk-tissue and/or single-cell data, and phenome-wide analyses. Results can be reviewed through an interactive dashboard, and the pipeline can run locally, on a high-performance computing cluster or in the cloud.

## 2 Methods

### 2.1 Implementation and inputs

drugMR is a Nextflow pipeline whose tasks run Python or R scripts that call the relevant statistical tools (Figure 1A). Inputs comprise one or more pQTL panels, an outcome GWAS and optional molecular QTL datasets listed in a manifest. Settings are specified in a YAML configuration file. The pipeline supports local, high-performance computing and cloud execution using Docker or Apptainer/Singularity containers. All settings below are configurable defaults.

**Figure 1:**
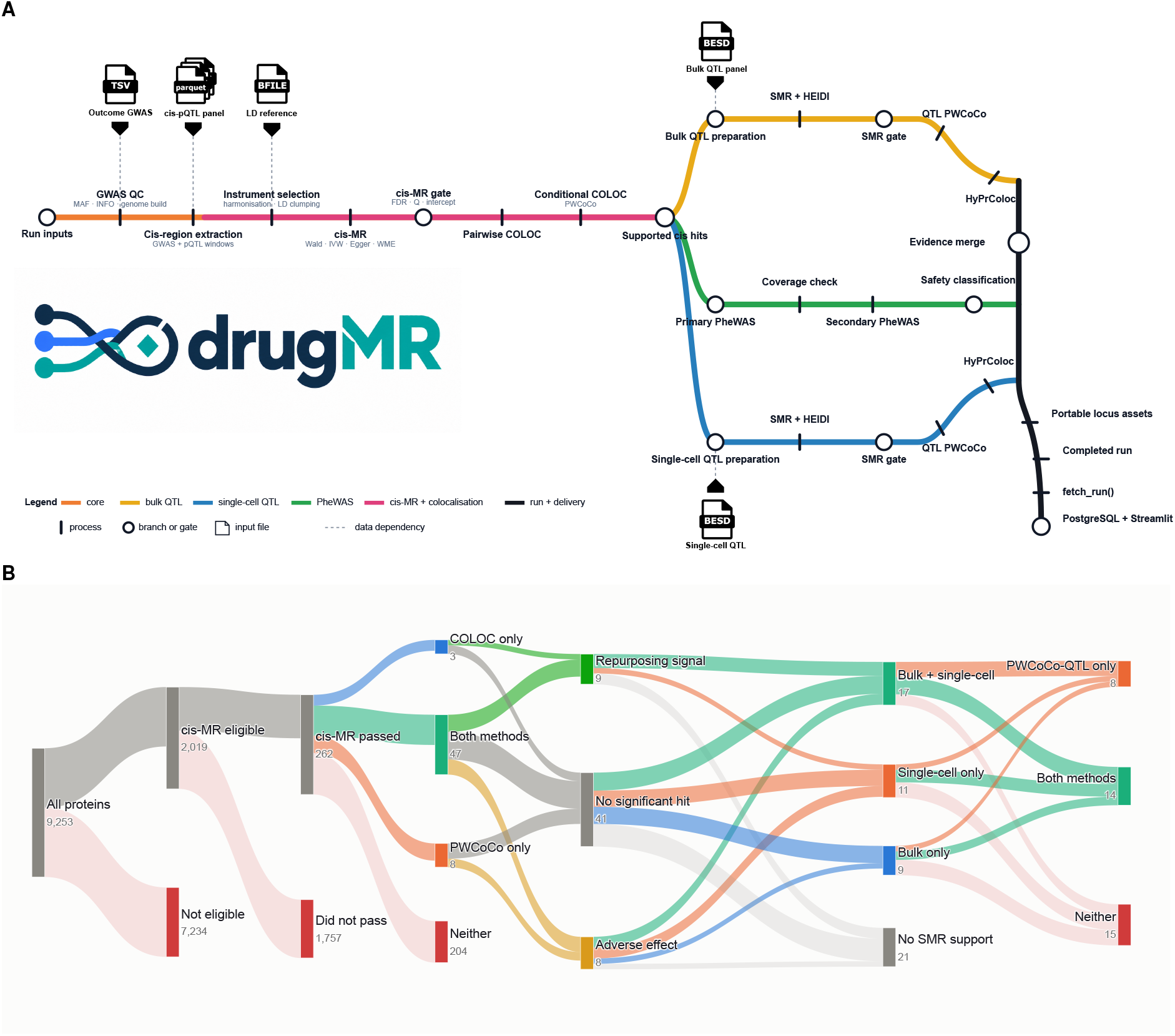
drugMR workflow and its application to SCZ. (A) Overview of the drugMR pipeline. Each line is a group of steps: GWAS quality control (orange), *cis*-MR and colocalisation (pink), bulk tissue QTL follow-up (yellow), single-cell QTL follow-up (blue), phenome-wide screening (green), and result delivery to the dashboard (black). Ticks mark individual steps, circles mark points where the workflow branches or applies a filter, and file icons mark input data. (B) Number of proteins retained at each stage of the SCZ analysis using the Wingo et al. (2025) brain pQTL panel. From left to right, columns show proteins with valid instruments, proteins passing *cis*-MR, colocalisation support by coloc, PWCoCo or both, phenome-wide screening results, SMR support in bulk or single-cell data, and three-trait colocalisation support by HyPrColoc, PWCoCo QTL or both. Phenome-wide results are shown for information and do not remove proteins.

### 2.2 Data preprocessing and quality control

The outcome GWAS is standardised and, where required, lifted over to the pQTL genome assembly (GRCh37/hg19 or GRCh38/hg38) using UCSC chain files and the Python liftover package (Hinrichs, 2006). Insertions and deletions, palindromic SNPs, duplicates and SNPs with minor allele frequency below 0.01 are removed where the required information is available. An imputation information score threshold may also be applied. The major histocompatibility complex region is excluded by default and the apolipoprotein E region optionally (see Supplementary table 1 for coordinates). For case-control GWAS, the effective sample size is 4*N*_cases_*N*_controls_*/*(*N*_cases_ + *N*_controls_). Proteins are mapped to their encoding genes using the NCBI Gene database (https://www.ncbi.nlm.nih.gov/datasets/gene/) for the chosen assembly. Each protein’s *cis* region is the window covered by its pQTL summary statistics, as supplied by the pQTL resource. Within this window, pQTL and GWAS variants are matched by SNP identifier. Linkage disequilibrium (LD) is estimated from a PLINK (Chang et al., 2015) format reference panel, by default the European ancestry samples of the 1000 Genomes Project Phase 3.

### 2.3 Instrument selection and *cis*-Mendelian randomisation

Instruments are selected for each protein according to the MR assumptions in Supplementary section 1.1. Eligible SNPs have *P <* 5 × 10^−8^ for protein abundance and *F* ≥ 10 (Burgess and Thompson, 2011). PLINK LD clumping keeps approximately independent instruments (*r*^2^ *<* 0.001 within 10 Mb). Effects are aligned to the pQTL risk allele and MR is run using TwoSampleMR (Hemani et al., 2018). The Steiger test compares the variance each instrument explains in protein abundance 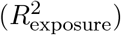 with that explained in the outcome 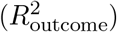 (Hemani et al., 2017). Instruments with 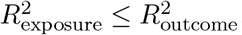, which suggest reverse causation, are removed by default. The primary estimate is the Wald ratio for one instrument (*n* = 1) and the inverse variance weighted (IVW) estimate for *n* ≥ 2 (Burgess et al., 2013). For *n* ≥ 3, MR Egger regression and the weighted median estimator are also run, and the MR Egger intercept is reported as a test for directional pleiotropy, without being used to exclude proteins. Primary *P* values are false discovery rate (FDR) corrected across all proteins in each pQTL panel. A protein passes *cis*-MR at *q*_FDR_ *<* 0.05 and, when *n* ≥ 2, it must also show no heterogeneity on Cochran’s Q test (*P*_*Q*_ *>* 0.05).

### 2.4 Pairwise colocalisation

Proteins passing *cis*-MR undergo colocalisation by two methods. coloc (Giambartolomei et al., 2014) assumes at most one causal variant per trait and uses *p*_1_ = 1 × 10^−4^, *p*_2_ = 1 × 10^−4^ and *p*_12_ = 1 × 10^−5^, with sensitivity analyses at *p*_12_ ∈ {5 × 10^−6^, 2 × 10^−6^, 1 × 10^−6^}. Because coloc may miss shared associations at loci containing multiple independent signals, Pair-Wise Conditional analysis and Colocalisation analysis (PWCoCo) (Robinson et al., 2022) is also applied. PWCoCo uses GCTA-COJO (Yang et al., 2012) to identify and condition independent signals before applying coloc to every protein-outcome signal pair and the unconditioned data. Either method supports colocalisation when any analysis has PP.H4 ≥ 0.70.

### 2.5 Molecular QTL follow-up

Genes encoding proteins that pass *cis*-MR and colocalisation are tested in bulk tissue or single-cell molecular QTL datasets, or both. Summary data based Mendelian randomisation (SMR) (Zhu et al., 2016) tests each molecular trait against the outcome, using its top QTL SNP (*P <* 5*×*10^−8^) as the instrument. The heterogeneity in dependent instruments (HEIDI) test, using SNPs with QTL *P <* 1.57 × 10^−3^, tests whether the association reflects a single causal variant shared by the molecular trait and the outcome, rather than distinct causal variants in LD. SMR *P* values are FDR corrected across all molecular traits in each dataset and tissue or cell type, and support requires *q*_SMR_ *<* 0.05 and *P*_HEIDI_ *>* 0.01.

Hypothesis prioritisation for multi-trait colocalisation (HyPrColoc) (Foley et al., 2021) tests whether protein abundance, the molecular trait and the outcome share one causal variant. The prior probability of a SNP being causal for one trait is 1 × 10^−4^ and the conditional prior is 0.02; sensitivity was assessed across prior.c ∈ {0.05, 0.02, 0.01, 0.005}. Support requires a posterior probability of at least 0.70 for a cluster of all three traits. The molecular QTL extension of PWCoCo (PWCoCo QTL) runs PWCoCo on three pairs: protein and outcome, molecular trait and protein, and molecular trait and outcome, with the last two using the same tissue or cell type. Support requires PP.H4 ≥ 0.70 in all three pairs, either for the same conditioned signal or in all three unconditioned analyses, in a tissue or cell type that also passed SMR and HEIDI.

### 2.6 Phenome-wide analyses

For each protein passing *cis*-MR and colocalisation, the drugMR PheWAS-MR module queries its instruments across ICD disease endpoints in FinnGen R13 (https://r13.finngen.fi/). Proteins without instruments in FinnGen are queried in the UK Biobank TOPMed-imputed PheWAS database (https://pheweb.org/UKB-TOPMed/). Effects are aligned to the protein-increasing allele and estimated using the Wald ratio for *n* = 1 or IVW for *n* ≥ 2. Bonferroni correction is applied per protein across 2,511 FinnGen endpoints (familywise *α* = 0.05). Significant associations are classified as possible repurposing opportunities when their direction agrees with the primary *cis*-MR estimate and as possible adverse effects when it is opposite.

### 2.7 Data storage and dashboard

Intermediate data are stored as Parquet files and final results as tab separated files. The drugmr Python package loads results into a PostgreSQL database, which serves a Streamlit dashboard. The dashboard shows each target’s MR, colocalisation, regional association, molecular QTL and phenome-wide results in one place; it is launched separately and is not a pipeline step.

## 3 Results

We applied drugMR to the PGC3 European schizophrenia (SCZ) GWAS (67,390 cases, 94,015 controls; Trubetskoy et al., 2022), using the Wingo et al. (2025) dorsolateral prefrontal cortex mass spectrometry pQTL panel (*N* = 1,013), with bulk brain (MetaBrain; *N* = 2,865) (de Klein et al., 2023) and single-cell (SingleBrain, seven cell types; *N* = 983) (Jang et al., 2026) eQTL data for molecular follow-up. All settings are listed in Supplementary section 1.2, and Figure 1B shows the targets retained at each stage.

Of 9,253 proteins in the panel, 2,019 had at least one valid instrument and were tested by MR (1,860 by Wald ratio, 159 by IVW). After FDR correction and the heterogeneity check, 262 were associated with SCZ (255 by Wald ratio, 7 by IVW; Supplementary table 2). Colocalisation supported 58 of these (PP.H4 ≥ 0.70; Supplementary table 3), which we refer to as proteogenomic targets. Coloc and PWCoCo agreed for 47, 3 were supported by coloc alone and 8 by PWCoCo alone (Supplementary table 4). Coloc support fell to 40, 23 and 13 proteins at *p*_12_ = 5 × 10^−6^, 2 × 10^−6^ and 1 × 10^−6^ (Supplementary table 3).

All 58 proteogenomic targets had instruments in FinnGen, so the UK Biobank fallback was not needed. Across 2,511 ICD disease endpoints, 8 had at least one Bonferroni significant association suggesting a potential adverse effect, 9 had only associations suggesting potential repurposing, and 41 had none (Supplementary table 5). PheWAS results are reported alongside each target and are not used to exclude any.

SMR and HEIDI supported 37 proteogenomic targets (17 in both bulk and single-cell data, 9 in bulk only, 11 in single-cell only; Supplementary table 6). HyPrColoc found colocalisation of protein abundance, gene expression and SCZ (posterior probability ≥ 0.70) for 14 of these: *ACADVL, ACTR1B, AK4, ANKRD45, CSDE1, CTPS1, DNAJC11, EEFSEC, ELAC2, IRF3, SLC25A24, STON2, TMX1* and *YOD1*. PWCoCo QTL supported all 14 and a further 8 (*ADSS2, CACNB3, CCDC92, KHK, MAPRE3, OGFOD2, PPP3CC* and *SPECC1*; Supplementary table 7). We refer to these 22 proteins as multi-omic targets. Four of them (*DNAJC11, EEFSEC, IRF3* and *OGFOD2*) also had a potential adverse effect association. Evidence for each target is in Supplementary table 8 and the dashboard.

The full workflow (16 tasks) ran as a single SLURM job on Supercomputing Wales (Falcon; 8 CPU cores, 64 GB of memory) in 4 h 19 min, using 25.4 CPU hours (Supplementary table 9; Supplementary Data 1, 2 and 3). The longest steps were *cis*-MR (1 h 17 min), single-cell and bulk SMR (56 and 49 min) and PWCoCo QTL (34 min). No task exceeded 7.3 GB of peak resident memory.

## 4 Conclusions and future perspectives

We developed drugMR, a Nextflow pipeline that prioritises drug targets by combining *cis*-MR, colocalisation, molecular QTL follow-up and phenome-wide screening in a single workflow. Previous proteome-wide MR studies have generally paired MR with single-signal colocalisation in bulk tissue (Zheng et al., 2020; Zhao et al., 2022). In the SCZ application, this approach would have missed 8 of 58 proteogenomic targets supported only by PWCoCo and 11 of 37 targets with SMR support found exclusively in single-cell data. Phenome-wide screening identified possible adverse-effect associations for 4 of the 22 multi-omic targets, providing information unavailable from MR and colocalisation alone. Because each dataset is listed in a manifest, the same analysis can be run with a different pQTL panel, outcome GWAS or molecular QTL dataset by editing the configuration file. The dashboard allows users without programming experience to review the evidence for each target. drugMR has two main limitations. First, its phenome-wide resources (FinnGen and UK Biobank) predominantly represent European ancestry, limiting generalisability to other populations. Second, it cannot yet test whether a protein affects disease through an intermediate biomarker. Future multivariable MR (Burgess and Thompson, 2015) and network MR (Burgess et al., 2015) subworkflows will address this, for example by testing whether a protein affects Alzheimer’s disease risk through amyloid-*β*42.

## Data availability

drugMR is available at https://github.com/guillermocomesanacimadevila/drugMR. Reference data and synthetic demo data are available at https://doi.org/10.5281/zenodo.22706705 and https://doi.org/10.5281/zenodo.22917384. SCZ GWAS summary statistics (Trubetskoy et al., 2022) were obtained from the PGC (https://pgc.unc.edu/for-researchers/download-results/), brain pQTLs from Wingo et al. (2025), bulk brain eQTLs from MetaBrain (https://www.metabrain.nl/) (de Klein et al., 2023), single-cell eQTLs (Jang et al., 2026) were sourced from Zenodo (https://zenodo.org/records/14908182). PheWAS summary statistics were queried from FinnGen release R13 portal (https://www.finngen.fi/en/access_results). SCZ results are provided in the Supplementary tables.

## References

N. Berdigaliyev and M. Aljofan. An overview of drug discovery and development. Future Medicinal Chemistry, 12(10), 2020. doi: 10.4155/fmc-2019-0307.

S. Burgess and S. G. Thompson. Avoiding bias from weak instruments in Mendelian randomization studies. International Journal of Epidemiology, 40(3):755–764, 2011. doi: 10.1093/ije/dyr036.

S. Burgess and S. G. Thompson. Multivariable Mendelian randomization: The use of pleiotropic genetic variants to estimate causal effects. American Journal of Epidemiology, 181(4):251–260, 2015. doi: 10.1093/aje/kwu283.

S. Burgess, A. Butterworth, and S. G. Thompson. Mendelian randomization analysis with multiple genetic variants using summarized data. Genetic Epidemiology, 37(7):658–665, 2013. doi: 10.1002/gepi.21758.

S. Burgess, R. M. Daniel, A. S. Butterworth, et al. Network Mendelian randomization: using genetic variants as instrumental variables to investigate mediation in causal pathways. International Journal of Epidemiology, 44(2):484–495, 2015. doi: 10.1093/ije/dyu176.

C. C. Chang, C. C. Chow, L. C. Tellier, et al. Second-generation PLINK: rising to the challenge of larger and richer datasets. GigaScience, 4(1), 2015. doi: 10.1186/s13742-015-0047-8.

N. M. Davies, M. V. Holmes, and G. Davey Smith. Reading Mendelian randomisation studies: a guide, glossary, and checklist for clinicians. BMJ, 362:k601, 2018. doi: 10.1136/bmj.k601.

J. A. de Amorim, J. V. F. Cavalcante, D. Marques-Coelho, et al. Causeway: a pipeline for genome-wide effector gene screening with Mendelian Randomization and colocalization. Bioinformatics Advances, 5(1), 2025. doi: 10.1093/bioadv/vbaf110.

N. de Klein, E. A. Tsai, M. Vochteloo, et al. Brain expression quantitative trait locus and network analyses reveal downstream effects and putative drivers for brain-related diseases. Nature Genetics, 55(3):377–388, 2023. doi: 10.1038/s41588-023-01300-6.

C. N. Foley, J. R. Staley, P. G. Breen, et al. A fast and efficient colocalization algorithm for identifying shared genetic risk factors across multiple traits. Nature Communications, 12(1), 2021. doi: 10.1038/s41467-020-20885-8.

C. Giambartolomei, D. Vukcevic, E. E. Schadt, et al. Bayesian test for colocalisation between pairs of genetic association studies using summary statistics. PLoS Genetics, 10(5):e1004383, 2014. doi: 10.1371/journal.pgen.1004383.

G. Hemani, K. Tilling, and G. Davey Smith. Orienting the causal relationship between imprecisely measured traits using GWAS summary data. PLOS Genetics, 13(11):e1007081, 2017. doi: 10.1371/journal.pgen.1007081.

G. Hemani, J. Zheng, B. Elsworth, et al. The MR-Base platform supports systematic causal inference across the human phenome. eLife, 7:e34408, 2018. doi: 10.7554/eLife.34408.

S. Hinrichs. The UCSC Genome Browser Database: update 2006. Nucleic Acids Research, 34 (Suppl_1):D590–D598, 2006. doi: 10.1093/nar/gkj144.

Jang, K. BP, A. Tokolyi, et al. A meta-analysis of single-nucleus expression quantitative trait loci linking genetic risk to brain disorders. Nature Genetics, 58(4):737–747, 2026. doi: 10.1038/s41588-026-02541-x.

E. V. Minikel, J. L. Painter, C. C. Dong, et al. Refining the impact of genetic evidence on clinical success. Nature, 629(8012):624–629, 2024. doi: 10.1038/s41586-024-07316-0.

S. Mundo, M. E. Grabowska, A. L. Dickson, et al. A Mendelian randomization-based drug repurposing pipeline with integrated AI-facilitated prioritization: application to lipid traits and coronary artery disease. medRxiv, 2026. doi: 10.64898/2026.02.28.26347341.

J. W. Robinson, G. Hemani, M. S. Babaei, et al. An efficient and robust tool for colocalisation: Pair-wise Conditional and Colocalisation (PWCoCo). bioRxiv, 2022. doi: 10.1101/2022.08.08.503158.

R. Santos, O. Ursu, A. Gaulton, et al. A comprehensive map of molecular drug targets. Nature Reviews Drug Discovery, 16(1):19–34, 2016. doi: 10.1038/nrd.2016.230.

A. F. Schmidt, C. Finan, M. Gordillo-Marañón, et al. Genetic drug target validation using Mendelian randomisation. Nature Communications, 11(1):3255, 2020. doi: 10.1038/s41467-020-16969-0.

D. Sun, W. Gao, H. Hu, et al. Why 90% of clinical drug development fails and how to improve it? Acta Pharmaceutica Sinica B, 12(7), 2022. doi: 10.1016/j.apsb.2022.02.002.

V. Trubetskoy, A. F. Pardiñas, T. Qi, et al. Mapping genomic loci implicates genes and synaptic biology in schizophrenia. Nature, 604(7906):502–508, 2022. doi: 10.1038/s41586-022-04434-5.

M. Verbanck, C. Y. Chen, B. Neale, et al. Detection of widespread horizontal pleiotropy in causal relationships inferred from Mendelian randomization between complex traits and diseases. Nature Genetics, 50(5):693–698, 2018. doi: 10.1038/s41588-018-0099-7.

A. P. Wingo, Y. Liu, S. M. Vattathil, et al. Multiancestry brain pQTL fine-mapping and integration with genome-wide association studies of 21 neurologic and psychiatric conditions. Nature Genetics, 57(9):2156–2165, 2025. doi: 10.1038/s41588-025-02291-2.

J. Yang, T. Ferreira, A. P. Morris, et al. Conditional and joint multiple-SNP analysis of GWAS summary statistics identifies additional variants influencing complex traits. Nature Genetics, 44(4):369–375, 2012. doi: 10.1038/ng.2213.

P. Yin and B. F. Voight. MeRP: a high-throughput pipeline for Mendelian randomization analysis. Bioinformatics, 31(6):957–959, 2015. doi: 10.1093/bioinformatics/btu742.

H. Zhao, H. Rasheed, T. H. Nøst, et al. Proteome-wide Mendelian randomization in global biobank meta-analysis reveals multi-ancestry drug targets for common diseases. Cell Genomics, 2(11):100195, 2022. doi: 10.1016/j.xgen.2022.100195.

J. Zheng, V. Haberland, D. Baird, et al. Phenome-wide Mendelian randomization mapping the influence of the plasma proteome on complex diseases. Nature Genetics, 52(10):1122–1131, 2020. doi: 10.1038/s41588-020-0682-6.

Z. Zhu, F. Zhang, H. Hu, et al. Integration of summary data from GWAS and eQTL studies predicts complex trait gene targets. Nature Genetics, 48(5):481–487, 2016. doi: 10.1038/ng.3538.

