## Supplementary_data for "drugMR: A multi-omics pipeline for genetically anchored drug-target discovery": Supp_data_1_execution_report_2026-09-26_00-20-44.html

[big\_plateau] Nextflow Workflow Report


Nextflow Report


- Summary
- Resources
- Tasks

[big\_plateau]

### Nextflow workflow report

#### `[big_plateau]`

Workflow execution completed successfully!

Run times
:   26-Sep-2026 00:20:45 - 26-Sep-2026 04:40:18
    (duration: **4h 19m 33s**)

16 succeeded

0 cached

0 ignored

0 failed (0 retries)

Nextflow command
:   ```
    nextflow run main.nf -profile slurm,local -params-file params/SCZ.wingo_brain.yaml --manifest_path assets/qtl_manifest.csv --container_bind /shared/scratch/SCWF00036/data
    ```

CPU-Hours
:   `25.4`

Launch directory
:   `/shared/scratch/SCWF00036/pipelines/drugMR`

Work directory
:   `/shared/scratch/SCWF00036/pipelines/drugMR/tmp/nf-work`

Project directory
:   `/shared/scratch/SCWF00036/pipelines/drugMR`

Script name
:   `main.nf`

Script ID
:   `2e0d8eb65e66fb7549aef709dfd7dbfb`

Workflow name
:   `drugMR`

Workflow version
:   `1.0.0`

Workflow description
:   drugMR: a multi-omics pipeline for genetically anchored drug-target discovery

Workflow home page
:   https://github.com/guillermocomesanacimadevila/drugMR

Workflow session
:   `01ec1fd9-77ff-4ca4-afd9-47f8288eb528`

Workflow profile
:   slurm,local

Nextflow version
:   version 26.04.0, build 12031 (29-04-2026 04:00 UTC)

## Resource Usage

These plots give an overview of the distribution of resource usage for each process.

#### CPU

- Raw Usage
- % Allocated

#### Memory

- Physical (RAM)
- Virtual (RAM + Disk swap)
- % RAM Allocated

#### Job Duration

- Raw Usage
- % Allocated

#### I/O

- Read
- Write

## Tasks

This table shows information about each task in the workflow. Use the search box on the right
to filter rows for specific values. Clicking headers will sort the table by that value and
scrolling side to side will reveal more columns.

Values shown as:

Human readable
Raw values

(tasks table omitted because the dataset is too big)

Generated by Nextflow, version 26.04.0
