## Supplementary_data for "drugMR: A multi-omics pipeline for genetically anchored drug-target discovery": Supplementary_material.pdf

### 1 Supplementary methods

#### 1.1 Assumptions of *cis*-Mendelian randomisation

Mendelian randomisation (MR) relies on three core assumptions: relevance, independence, and exclusion restriction (Supplementary Fig. 1). Here, these assumptions are described in the context of a *cis*-MR analysis evaluating the effect of genetically proxied protein abundance, denoted by $X$, on Schizophrenia (SCZ) risk, denoted by $Y$.

**Relevance assumption.** The genetic instruments of use, denoted as $G_j$, must be strongly associated with the exposure phenotype $X$ (in this case, the abundance of protein $X$ measured in a given pQTL panel). This is commonly assessed using genome-wide significant exposure associations, for instance $P_{jX} < 5 \times 10^{-8}$, and sufficient instrument strength, commonly $F_{jX} > 10$. For example, in a protein $X \rightarrow$ SCZ *cis*-MR analysis, SNPs associated with protein $X$ at genome-wide significance and passing the $F_{jX}$ threshold would satisfy this assumption.

**Independence assumption.** The genetic instruments of use, $G_j$, must be independent of confounders, denoted as $U$, affecting both protein abundance, $X$, and SCZ risk, $Y$. For example, in a protein $X \rightarrow$ SCZ *cis*-MR analysis, instruments associated with smoking, adiposity, systemic inflammation, or other factors influencing both protein abundance and SCZ risk through pathways independent of protein $X$ could violate the independence assumption. An association between $G_j$ and $U$ is denoted as $\phi_j$.

**Exclusion restriction.** The genetic instruments, $G_j$, must influence SCZ risk, $Y$, only through protein abundance, $X$, and not via alternative biological pathways. For example, in a protein $X \rightarrow$ SCZ *cis*-MR analysis, a *cis*-pQTL could violate this assumption if it independently affected a neighbouring gene or another protein that influenced SCZ risk through a pathway unrelated to protein $X$. Violations of this assumption reflect horizontal pleiotropy, denoted as $\alpha_j$.

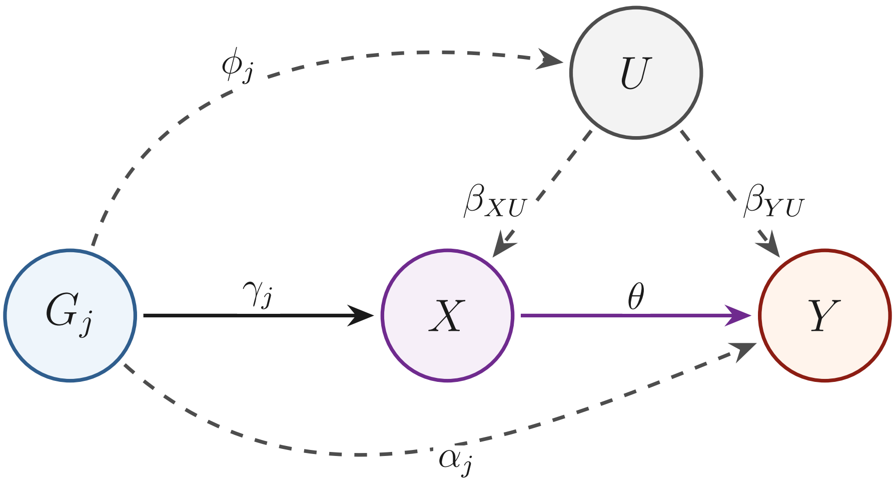


Figure 1: **Core assumptions underlying Mendelian randomisation.** Directed acyclic graph illustrating the three core assumptions required for valid MR inference in a protein $X \rightarrow$ schizophrenia analysis. $G_j$ represents the genetic instrument, $X$ the abundance of the instrumented protein, $Y$ SCZ risk, and $U$ an unmeasured confounder affecting both $X$ and $Y$. The causal effect of protein abundance on SCZ risk is denoted by $\theta$. The relevance assumption requires the genetic instrument to be associated with protein abundance, denoted by $\gamma_j$. The independence assumption requires the genetic instrument to be independent of $U$, such that $\phi_j \approx 0$. The exclusion restriction assumption requires the instrument to influence $Y$ only through $X$, such that $\alpha_j \approx 0$. Dashed arrows represent potential violations of the independence and exclusion restriction assumptions.

#### 1.2 Schizophrenia analysis configuration

The complete configuration used to apply drugMR to the SCZ genome-wide association study (GWAS) using a brain pQTL dataset is reproduced below. Downstream molecular QTL triangulation used MetaBrain bulk-brain eQTLs and SingleBrain single-nucleus eQTLs across seven major brain cell types: astrocytes, endothelial cells, excitatory neurons, inhibitory neurons, microglia, oligodendrocytes, and oligodendrocyte precursor cells. Parameter definitions and additional usage instructions are available in the drugMR documentation.


```
pheno_id: SCZ
sumstats: ../../data/GWAS/SCZ.tsv
n_cases: 67390
n_controls: 94015
genome_build: GRCh37
target_build: GRCh38

snp_col: SNP
a1_col: A1
a2_col: A2
beta_col: BETA
se_col: SE
p_col: P
pos_col: POS
chr_col: CHR
af_col: FRQ

pqtl_dataset: wingo_brain

reference_data:
  ref_bfile: ref/1000G_EUR_Phase3_plink/1000G.EUR.QC.ALL
  liftover_dir: ref/liftover
  gene_annotation: ref/NCBI/NCBI_genes_grch38_with_synonyms.tsv

run_smr: true
bulk_qtl_datasets: [metabrain]
sc_qtl_dataset: singlebrain

maf: 0.01
overwrite: false
remove_mhc: true
remove_apoe: false

gates:
  cis_mr:
```

```yaml
  wald_fdr_q: 0.05
  ivw_fdr_q: 0.05
  cochran_q_pval: 0.05
  egger_intercept_pval_min: 0
  min_instruments_for_ivw: 3
  apply_steiger_filter: true
  clump_kb: 10000
  clump_r2: 0.001
  instrument_pval_threshold: 5.0e-8
  min_f_stat: 10

coloc:
  pp4_threshold: 0.7
  p1: 1.0e-4
  p2: 1.0e-4
  p12: 1.0e-5

smr:
  p_qtl_smr: 5.0e-8
  p_qtl_heidi: 1.57e-3
  diff_freq_prop: 0.55
  p_smr_threshold: 0.05
  p_heidi_threshold: 0.01

hyprcoloc:
  prior_1: 1.0e-4
  prior_c: [0.05, 0.02, 0.01, 0.005]
  reg_thresh: [0.5, 0.6, 0.7]
  align_thresh: [0.5, 0.6, 0.7]
  equal_thresholds: true

pwcoco:
  pp4_threshold: 0.7

phewas:
  bonferroni_alpha: 0.05
  coloc_threshold: 0
```
